# Curvature effect of PE-included membrane on the behavior of cinnamycin on the membrane

**DOI:** 10.1101/2020.06.19.161679

**Authors:** S-R. Lee, Y. Park, J-W. Park

## Abstract

The behavior of the cinnamycin on the biomimetic membrane was studied with respect to the curvature of the phosphatidylethanolamine(PE)-included membrane with the adhesion measured by the atomic force microscope(AFM). The membrane was formed through vesicle fusion on the hydrophobic surface of the sphere spheres, which was used to define the curvature of the membrane. The hydrophobicity was generated by the reaction of alkyl-silane and analyzed with the X-ray photoelectron spectrometer. The cinnamycin, immobilized covalently to the AFM tip coated with 1-mercapto-1-undecanol that was observed inert to any adhesion to the membrane, showed that the adhesion became stronger with the increase in the curvature. The correlation between the adhesion and the curvature was linearly proportional. Previously, it was found that the cinnamycin was bound to PE headgroup and the binding was enhanced by the interaction of the hydrophobic area located at one side of the cinnamycin. Therefore, the linear proportionality seems to suggest that the interaction is related to the one dimensional orientation of the binding.

**Statement of Significance:** The behavior of the cinnamycin was studied on the phosphatidylethanolamine(PE)-included membrane with respect to the curvature of the membrane. The cinnamycin, immobilized covalently to the atomic force microscope, showed that the adhesion became stronger linearly with the increase in the curvature. Previously, it was found that the specific binding between the cinnamycin and PE headgroup was enhanced by the interaction of the hydrophobic area located at one side of the cinnamycin. Therefore, the linear proportionality seems to suggest that the interaction is related to the one dimensional orientation of the binding.

## Introduction

Cinnamycin is a 19-amino acid tetracyclic and electrically neutral peptide capable of forming an equimolecular complex through the specific binding with phosphatidylethanolamine(PE) (1,2). This peptide has been focused on to investigate PE-involved processes such as apoptosis, cell division, migration, and tumor vasculature, since the unique binding was found (3-7). In eukaryotic cells, PE is distributed in the inner layer of the plasma membrane (8-10). It has been shown that cinnamycin led phospholipid movement across the bilayer in a PE-dependent manner (11). This movement seemed to induce the inner layer PE to the peptide and promote the binding of the cinnamycin. With high PE-concentration, it was observed that the membranes were reorganized and their structures were altered. However, the detailed information of the mechanism is little known.

Supported lipid layers have been employed for bio-mimetic membrane study due to several advantages such as ease of formation, convenience of control over complexity, assurance of stability, and capability of utilizing a wide range of different techniques for highly sensitive surface analysis (12). They have been used in many areas such as active transport, energy production, and signal transduction in layer-associated systems (13). The various types of the layers been characterized with numerous surface analytical techniques, including atomic force microscopy, surface plasmon resonance spectrometry, surface-enhanced Raman spectrometry, reflection-absorption infrared spectrometry, total internal reflection fluorescence microscopy, cyclic voltammetry, and the quartz crystal resonance (14-17). The atomic force microscope(AFM) has been extensively popular for measuring intermolecular forces due to its pico-newton force sensitivity and nanometer positional accuracy. Furthermore, the AFM has been capable of operation under physiological conditions (18,19). The force sensitivity has made the AFM available to magnitude of molecular recognition that may be generated from non-covalent bonds – hydrogen, hydrophobic, ionic, and van der Waals interactions (20). In this work, we aim to investigate the quantitative curvature effect of the PE-included membrane on the direct behavior of cinnamycin on biomimetic membranes.

## Experiments

Lipid layer was formed with two steps. First, a hydrophobic silica-nanosphere surface was prepared with silica nanospheres and alkane silane chemistry. As the second step, the lipid layer was assembled on the hydrophobic surface through vesicle fusion by hydrophobic interaction. For the detailed first-step, aqueous silica-sphere solutions was changed into a methanolic solution through a gradual increase of methanol ratio in the solution for facile solvent-evaporation without sphere aggregation. The sphere solutions with a desired diameter were purchased from Polyscience, Inc. (Warrington, PA) and Bangs Laboratories, Inc. (Fishers, IN) in dispersion of 5% in aqueous solution. Several drops of the methanolic solution were spread on each piece of a silicon wafer (Wafer Market Co., Ltd., Seongnam-si, Gyeonggi-do, Republic of Korea), cut by 1 cm by 1 cm. After the methanol evaporation, the spheres were transferred to ultra-violet cleaner with ozone treatment (UVO Cleaner Model 18, Jelight, Irvine, CA). Then, the spheres were covered carefully with silane monolayer using octadecyltrimethoxysilane (OTS) (Sigma, St. Louis, MO) solution (10/90, v/v, OTS/toluene) under nitrogen condition at 80 °C for 2 h, as described previously (21). The wafer was transferred in a bath-type sonicator to remove unreacted OTS molecules. After the sonication, the elements of the support surfaces were analyzed with X-ray photoelectron spectrometer (XPS) (Sigma Probe, Thermo VG Scientific, West Sussex, UK). The structures of the supports were confirmed with scanning electron microscope (SEM) (JSM-6700F, Jeol Ltd., Tokyo, Japan). The second step for the lipid-layer formation was performed in the vesicle preparation and the AFM experiments described later.

For the vesicle preparation, dioleoylphosphatidylcholine(DOPC) and dioleoyl-phosphatidylethanolamine(DOPE) from Avanti (Alabaster, AL) were dissolved at 90:10 ratio (DOPC:DOPE) or pure DOPC in chloroform. And the chloroform was subsequently evaporated under a dry stream of nitrogen to form the lipid films at the wall of a glass tube. The lipid films were at low pressure for several hours to remove last traces of the solvent and immersed overnight at room temperature in 2 ml of the solution containing 10 mM PBS at pH 7 for hydration. The hydrated solution was through freezing and thawing with the vigorous vortex every 10 minute ten cycles, and through extrusion of two stacked 100 nm pore size polycarbonate filters at room temperature to achieve the formation of uni-lamellar vesicles. The vesicle solution was transferred to the instrument of dynamic light scattering (ELS-8000, Otsuka, Tokyo, Japan) to measure the diameter of the vesicles, which was distributed normally between 130 and 170 nm.

AFM measurements were performed at room temperature with the sensitivity of ∼0.01 nN (Nanoscope v5.12, Veeco, Santa Barbara, CA) (22). The cinnamycin (Sigma) was immobilized on gold-coated cantilevers (OMCL-TR400PB, Olympus, Tokyo, Japan) with the overnight-immersion in an ethanoic solution containing 1 mM of 90% 1-mercapto-1-undecanol(MUD) and 10% 16-mercaptohexadecanoic acid or 100% MUD, the 30 minute treatment with a solution containing 25 mg/ml 1-ethyl-3-(3-dimethylaminopropyl)-carbodiimide and 10 mg/ml *N*-hydroxysuccinimide, and the 1 hour incubation with 0.1 mg/ml cinnamycin. After each step, the cantilever was washed out with the PBS solution thoroughly. The cinnamycin-immobilized-cantilever was stored in PBS solution right before it was used for the measurements, in order to avoid any dewetting on the tip of the cantilever. All chemicals essential for the immobilization were acquired from Sigma. The hydrophobic silica-nanosphere surface was placed on the AFM scanner, and approached to the cantilever tip loaded onto a liquid cell. The cell-inside, sealed completely with a silicon O-ring, was fully filled with the vesicle solution. The hydrophobic surface inside the cell was exposed to the solution with waiting 1 hour at room temperature for the vesicle fusion to form lipid layer on the surface. After remained vesicles were washed out by circulating the PBS solution inside of the cell, the retracted deflection curves were collected with the velocity of 20 nm/s. After more than 200 curves were collected, only the curves with the highest-adhesion-point among them were selected. The deflection curves were converted with 0.09 N/m spring constant provided from the manufacturer into the force curves, which were further analyzed.

## Results and Discussion

Toward the goal of investigating curvature effect of PE-included membrane on the interaction of cinnamycin, a phospholipid layer on the hydrophobic surface generated from OTS treatment were assembled through liposome fusion. Fig. 1 shows the schematic structure of biomimetic membrane, which was intended to be achieved. First, four different sizes of silica spheres were uniformly distributed on each silicon wafer prior to OTS treatment. After the treatment, the OTS monolayer was characterized using XPS. The results of the XPS analysis were almost identical with the study published previously (23). The results suggested that OTS-treated surface contained large amounts of silicon, oxygen, and carbon. The carbon has a C1s 285.6 eV binding energy associated with OTS. The relative abundance and binding energies of elements associated with silicon dioxide and the treatment presents that adsorption of OTS leads to a layer of irreversibly bound octadecyl groups on the surface of the silica spheres deposited on silicon wafer. The surface of the sphere arrays were shown in Fig. 2.

**Figure. 1.**
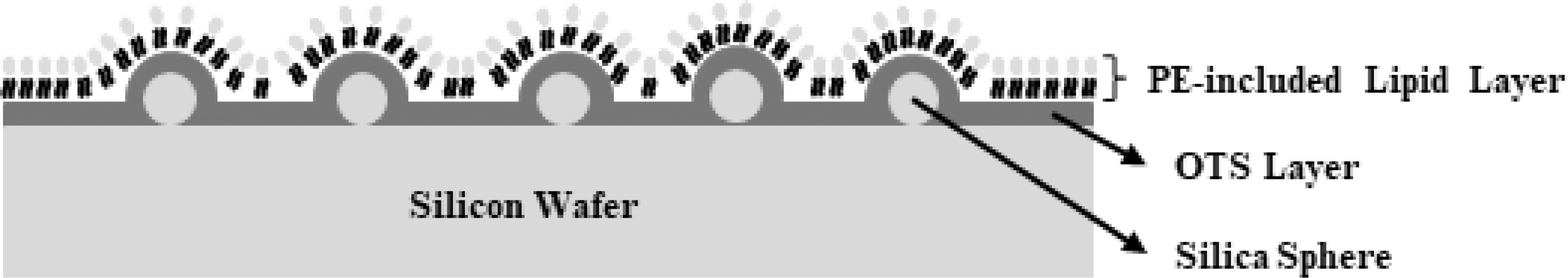
Schematic drawing of the PE-included lipid layers formed the hydrophobic surfaces with different curvatures.

**Figure 2.**
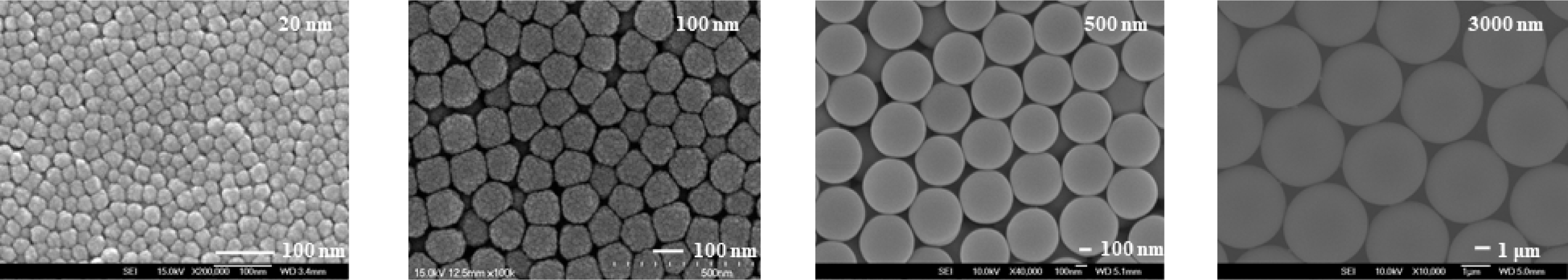
Silica spheres deposited on silicon wafer, which define the curvature of the PE-included lipid layers.

Deflection measurements were performed at room temperature on the lipid layer formed on the hydrophobic surfaces deposited on the silica spheres. The force between the MUDs on the tip and the lipid layer on the 20 nm-sphere structure, converted from the deflection, is plotted with respect to the distance between them (Fig. 3). The plot includes tiny forces in the long-range (> 6nm). At the buffer solution, the theoretical force, Derjaguin-Landau-Verwey-Overbeek force, was calculated for comparison. The force is the sum of the double-layer force and the van der Waals force (Hamaker constant 7.0×10^−21^ J for hydrocarbon) and lack of the long-range as well (24). However, the plot shows a net repulsive force at short-range, which may originate from steric/hydration on the surface (25). Since any significant attractive force is absent, the MUD layer is an excellent system to monitor a specific binding of a protein to supported lipid layers. The behaviors of the MUDs were also found identically at the other diameters of the surface.

**Figure 3.**
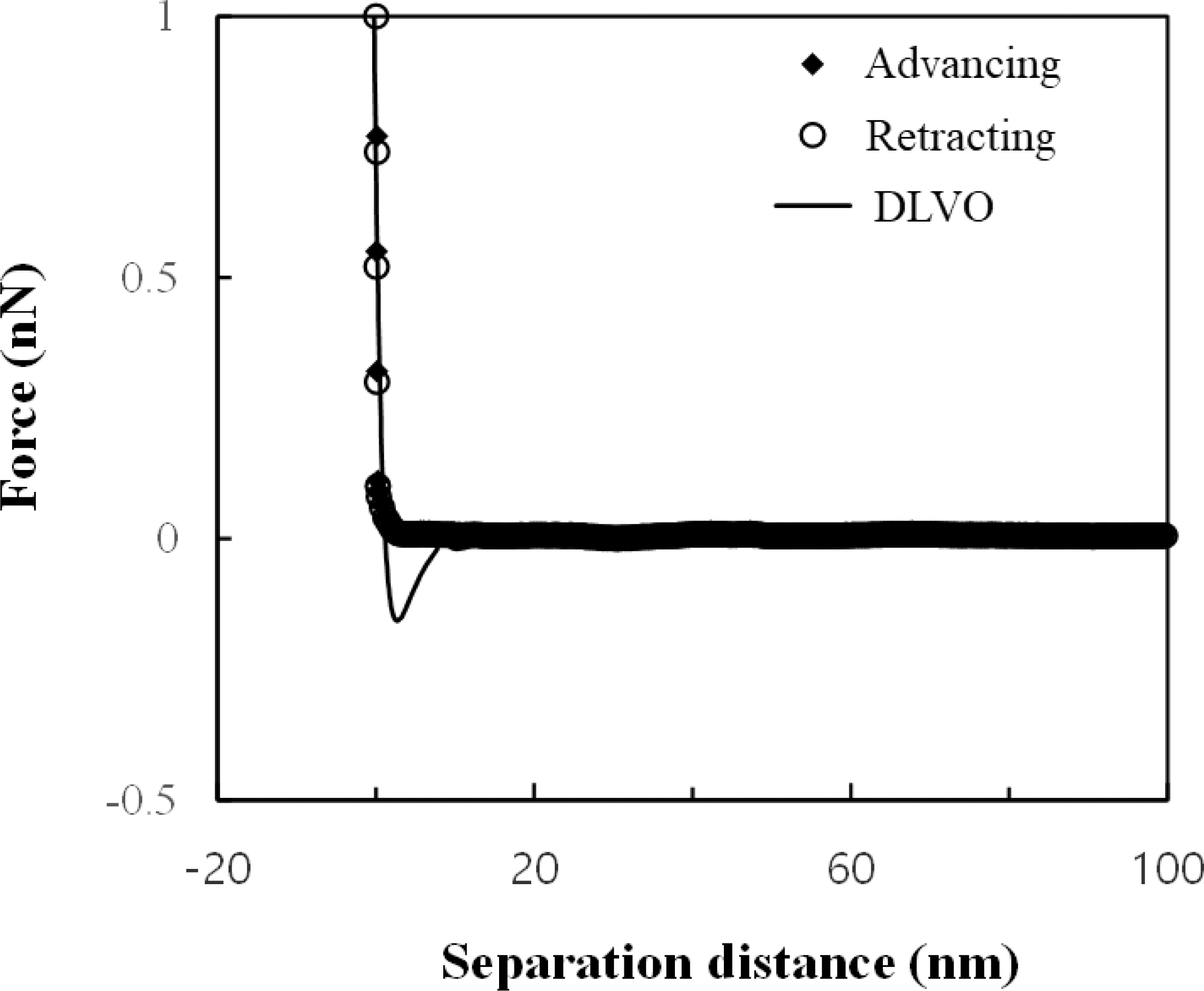
Forces between the PE-included lipid layer and MUDs immobilized on the tip.

The force between the pure DOPC outer-layer on the 20 nm-sphere structure and the cinnamycin on the tip was measured with respect to the distance between them (Fig. 4, solid line). The force included the adhesive between 0.06 and 0.1 nN, which was different from that between the MUDs and the lipid layer. The cinnamycin was believed to cause the adhesion that may be interpreted into a change in the magnitude of the short-range repulsive force or/and the van der Waals attractive force. In addition, the force on the DOPE-included outer-layer also showed adhesion. However, the adhesive force was even more dominant than that of the pure DOPC outer-layer (Fig. 4, dash line). At 10% DOPE, the force was between 0.55 and 0.65 nN with the range of ±0.02 nN for each point. Furthermore, the distinct snap-off, the abrupt break of the adhesion, was found at the layer including DOPE, while it was little shown without DOPE. Therefore, the force measured on the pure DOPC outer-layer seems non-specificity and the significant increase in the adhesive force was apparently generated specifically by DOPE and the cinnamycin.

**Figure 4.**
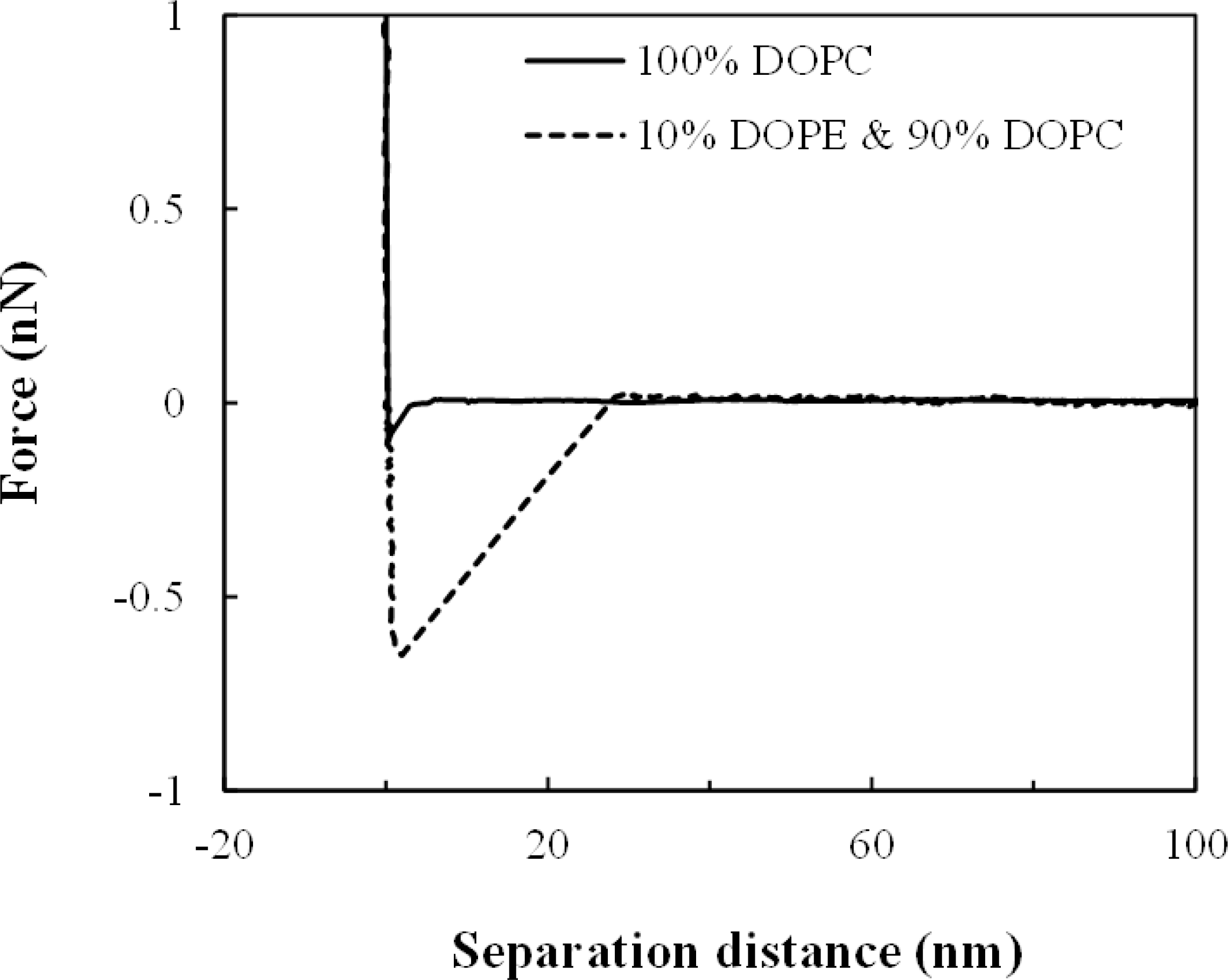
Force between the PE-included lipid layer and the cinnamycins immobilized on MUD-coated the tip.

The substantial adhesive interaction may be caused by either the dissociation of the DOPE-cinnamycin specific-binding or the dislocation of DOPE from the lipid layer. This confusion was clarified with two observations. The repetition of the identical adhesions was found between the DOPE lipid layer and the cinnamycin on the tip. The repetition indicated that the layer was recovered in short time right after the modification caused by the measurements. Second, in the previous AFM studies, it was found that the supported lipid layer was unperturbed with forces even more than 1.0 nN (26). Therefore, the binding between DOPE and cinnamycin was dissociated without any displacement of the lipids in the layer. The number of the cinnamycin-DOPE bindings was evaluated from the estimation of the contact area between the tip and the lipid layer (27). For the tip of 20 nm radius used for the measurement, the contact area on the lipid layer on the 20 nm-sphere structure was about 175 nm^2^ under a 1 nN load (28). Considering the radius of the cynnamycin gyration, about 8 interactions seemed to occur in the area for the specific binding. Besides the number of the interactions, the variations of the DOPE-orientation in the layer and the binding-site location deep within the cinnamycin might affect on the change in the adhesion magnitude at each condition.

The difference in the significant adhesion was also observed with respect to the curvature of the lipid layer. Prior to the measurements, it was expected that the increase in the curvature led to the decrease in the magnitude of the force. This expectation was induced from the increase of the contact area where the force acted, because the area was 175 to 260 nm^2^ from 20 nm to 3 µm-sphere structure. However, it was found that the force was correlated proportionally with the curvature. The adhesive force was 0.55∼0.65 nN for the lipid layer on the 20 nm-sphere structure, while the force was 0.4∼0.5 nN for the layer on the 3 µm-sphere structure. The range of the force was compared with that for the lipid layer on the planar surface. The ratio of the lipid thickness to 20 nm diameter was even more than 0.1, while the ratio for 3 µm sphere was much less than 0.001. The previous study suggested that the force for the lipid layer on the 3 µm-sphere was almost identical with that on the planar surface. (29).

According to the curvature of the PE-included membrane, the adhesive forces were changed. The force at each curvature was summarized in Table 2. From 20 to 3000 nm, the adhesive force was gradually decreased. Since the geometric exposure of a single PE to the environment was increased proportionally with square of the radius, the decrease in the adhesive force was considered as a function of relevance to curvature square. However, it was found that the decrease was proportional linearly to the curvature. Previously, it was found that the cinnamycin was bound to PE headgroup and the binding was enhanced by the hydrophobic interaction (2). And the interaction is inferred to occur at one side of the structure since the cinnamycin has an amphiphilic structure (30). Therefore, the linear relation seems to suggest that the interaction is related to the one dimensional orientation of the binding.

## Conclusion

In this study, the effect of the PE-included membrane curvature was investigated on the interaction between the cinnamycin and the membrane with the AFM adhesion measurements. The membrane was formed through vesicle fusion on the hydrophobic surface of the silica spheres used to define the curvatures. The cinnamycin was immobilized covalently to the cantilever tip coated with MUD that was without any adhesion to the membrane, and showed the behavior of the stronger adhesion with the increase in the curvature. The strength change was linearly proportional to the curvature, not the curvature square that the geometric exposure of a single PE was a function of. In the previous study, it was found that the cinnamycin was bound to PE headgroup and the binding was enhanced by the interaction of the hydrophobic area located at one side of the cinnamycin. Therefore, the linear proportionality seems to suggest that the interaction is related to the one dimensional orientation of the binding.

## Author Contributions

Jin-Won conceived, supervised the study, and designed experiments; Sang-Ryong and Yeseul performed experiments; Jin-Won and Sang-Ryong analysed data and wrote the manuscript.

## Acknowledgments

This study was supported by the Research Program funded by the SeoulTech (Seoul National University of Science and Technology).

## Tables

**Table 1.**
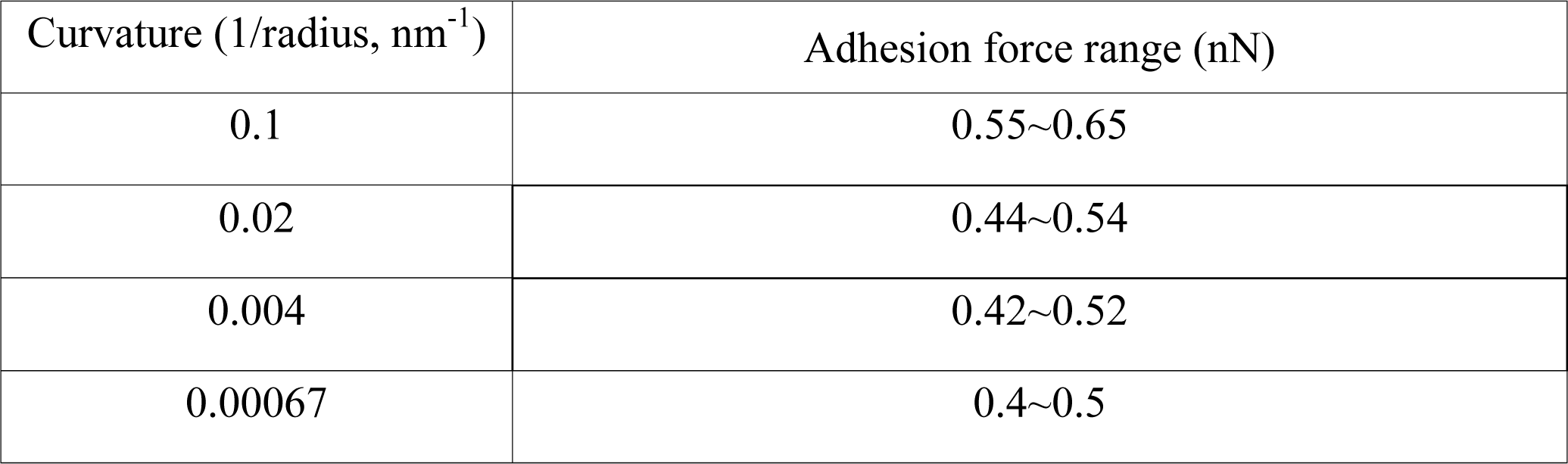
Adhesion forces up on the curvature of the PE-included membrane

| Curvature (1/radius, nm<sup>-1</sup>) | Adhesion force range (nN) |
| --- | --- |
| 0.1 | 0.55~0.65 |
| 0.02 | 0.44~0.54 |
| 0.004 | 0.42~0.52 |
| 0.00067 | 0.4~0.5 |

## References

1. Wakamatsu, K., S.Y. Choung, T. Kobayashi, K. Inoue, T. Higashijima, and T. Miyazawa. 1990. Complex formation of peptide antibiotic Ro09-0198 with lysophosphatidylethanolamine: 1H NMR analyses in dimethyl sulfoxide solution. Biochemistry 29:113–118.

2. Machaidze, G., A. Ziegler, and J. Seelig. 2002. Specific binding of Ro 09-0198 (cinnamycin) to phosphatidylethanolamine: a thermodynamic analysis. Biochemistry 41:1965–1971.

3. Emoto, K., H. Inadome, Y. Kanaho, S. Narumiya, and M. Umeda. 2005. Local change in phospholipid composition at the cleavage furrow is essential for completion of cytokinesis. J. Biol. Chem. 280:37901–37907.

4. Kato, U., H. Inadome, M. Yamamoto, K. Emoto, T. Kobayashi, and M. Umeda. 2013. Role for phospholipid flippase complex of ATP8A1 and CDC50A proteins in cell migration. J. Biol. Chem. 288:4922–4934.

5. Tafesse, F.G., A.M. Vacaru, E.F. Bosma, M. Hermansson, A. Jain, A. Hilderink, P. Somerharju, and J.C.M. Holthuis. 2014. Sphingomyelin synthase-related protein SMSr is a suppressor of ceramide-induced mitochondrial apoptosis. J. Cell Sci. 127:445–454.

6. Stafford, J.H., and P.E. Thorpe. 2011. Increased Exposure of Phosphatidylethanolamine on the Surface of Tumor Vascular Endothelium. Neoplasia 13:299–308.

7. Phoenix, D.A., F. Harris, M. Mura, and S.R. Dennison. 2015. The increasing role of phosphatidylethanolamine as a lipid receptor in the action of host defence peptides. Prog. Lipid Res. 59:26–37.

8. Devaux, P.F. 1991. Static and dynamic lipid asymmetry in cell membranes. Biochemistry 30:1163–1173.

9. Cerbon, J., and V. Calderon. 1991. Changes of the compositional asymmetry of phospholipids associated to the increment in the membrane surface potential. Biochim. Biophys. Acta. 1067:139–144.

10. Zachowski, A. 1993. Phospholipids in animal eukaryotic membranes: transverse asymmetry and movement. Biochem. J. 294:1–14.

11. Makino, A., T. Baba, K. Fujimoto, K. Iwamoto, Y. Yano, N. Terada, S. Ohno, S.B. Sato, A. Ohta, M. Umeda, K. Matsuzaki, and T. Kobayashi. 2003. Cinnamycin (Ro 09–0198) promotes cell binding and toxicity by inducing transbilayer lipid movement. J. Biol. Chem. 278:3204–3209.

12. Lee, S.-R., and J.-W. Park. 2018. Trehalose-Induced Variation in Physical Properties of Fluidic Lipid Bilayer. J. Membr. Biol. 251:705–709.

13. Gennis, R.B. 1989. Biomembranes: molecular structure and function. Springer, New York.

14. Meuse, C.W., S. Krueger, C.F. Majkrzak, J.A. Dura, J. Fu, J.T. Connor, and A.L. Plant. 1998. Hybrid bilayer membranes in air and water: infrared spectroscopy and neutron reflectivity studies. Biophys. J. 74:1388–1398.

15. Ohlsson, P.A., T. Tjärnhage,E. Herbai, S. Löfås, and G. Puu. 1995. Liposome and proteoliposome fusion onto solid substrates, studied using atomic force microscopy, quartz crystal microbalance and surface plasmon resonance. Biological activities of incorporated components. Bioelectrochem. Bioenerg. 38:137–148.

16. Schmidt, E.K., T. Liebermann, M. Kreiter, A. Jonczyk, R. Naumann, A. Offenhausser, E. Neumann, A. Kukol, A. Maelicke, and W. Knoll. 1998. Incorporation of the acetylcholine receptor dimer from Torpedo californica in a peptide supported lipid membrane investigated by surface plasmon and fluorescence spectroscopy. Biosens. Bioelectron. 13:585–591.

17. Terrettaz, S., T. Stora, C. Duschl, and H. Vogel. 1993. Protein binding to supported lipid membranes: investigation of the cholera toxin-ganglioside interaction by simultaneous impedance spectroscopy and surface plasmon resonance. Langmuir 9:1361–1369.

18. Liedberg, B., C. Nylander, and I. Lundström. 1983. Surface plasmon resonance for gas detection and biosensing. Sensors & Actuators 4:299–304.

19. Yuan, J., R. Oliver, J. Li, J. Lee, M. Aguilar, and Y. Wu. 2007. Sensitivity enhancement of SPR assay of progesterone based on mixed self-assembled monolayers using nanogold particles. Biosens. Bioelectron. 23:144–148.

20. Grieshaber, D., R. MacKenzie, J. Vörös, and E. Reimhult. 2008. Electrochemical Biosensors - Sensor Principles and Architectures. Sensors 8:1400–1458.

21. Maoz, R., and J. Sagiv. 1984. On the formation and structure of self-assembling monolayers. I. A comparative atr-wettability study of Langmuir—Blodgett and adsorbed films on flat substrates and glass microbeads. J. Colloid Interface Sci. 100:465–496.

22. Putman, C.A.J., B.G. de Grooth, N.F. van Hulst, and J. Greve. 1992. A detailed analysis of the optical beam deflection technique for use in atomic force microscopy. J. Appl. Phys. 72:6–12.

23. Jones, R.L., B.L. Harrod, and J.D. Batteas. 2010. Intercalation of 3-phenyl-1-proponal into OTS SAMs on silica nanoasperities to create self-repairing interfaces for MEMS lubrication. Langmuir 26:16355–16361.

24. Israelachivili, J.N. 1991. Intermolecular & Surface Forces. Academic Press, New York.

25. Ducker, W.A., T.J. Senden, and R.M. Pashley. 1991. Direct measurement of colloidal forces using an atomic force microscope. Nature 359:239–241.

26. Park, J.-W., and D.J. Ahn. 2008. Temperature effect on nanometer-scale physical properties of mixed phospholipid monolayers. Colloid Surf. B: Biointerfaces 62:157–161.

27. Laney, D.E., R.A. Garcia, S.M. Parsons, and H.G. Hansma. 1997. Changes in the Elastic Properties of Cholinergic Synaptic Vesicles as Measured by Atomic Force Microscopy. Biophys. J. 72:806–813.

28. Park, J.-W. 2010. Probe chemistry effect on surface properties of asymmetric-phase lipid bilayers. Colloid Surf. B: Biointerfaces 75:290–293.

29. Kim, S.-E., and J.-W. Park. 2020. Analysis of interactions between cinnamycin and biomimetic membranes. Colloids Surf. B: Biointerfaces 185:110595.

30. Choung, S.Y., T. Kobayashi, K. Takemoto, H. Ishitsuka, and K. Inoue. 1988. Interaction of a cyclic peptide, Ro09-0198, with phosphatidylethanolamine in liposomal membranes. Biochim. Biophys. Acta 940:180–187.

